## Supplementary material for "Coordinated regulation of chemotaxis and resistance to copper by CsoR in *Pseudomonas putida*": Fig. S1; Fig. S2; Fig. S3; Fig. S4; Fig. S5; Fig. S6; Fig. S7; Table S1; Table S2; Table S3.

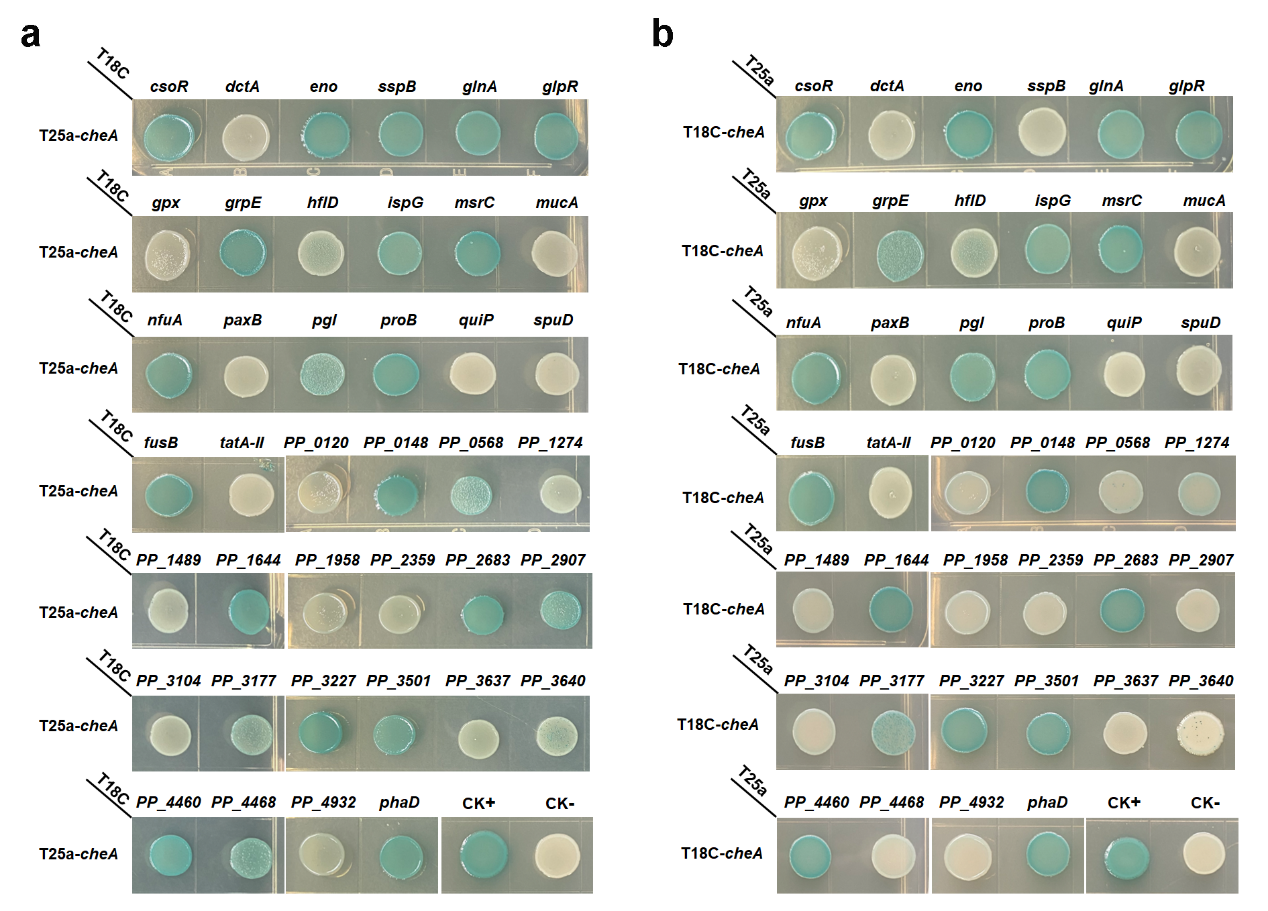

**Fig. S1** Detect the interaction between CheA and the 40 proteins using BTH. CheA was cloned into T25a (a) and T18C (b). Blue indicates protein-protein interaction, and white indicates no interaction in the colony after 60 h of incubation. A colony containing T25a-zip and T18C-zip plasmids was used as a positive control (CK+), and a colony containing empty T25a and T18C plasmids was used as a negative control (CK-).

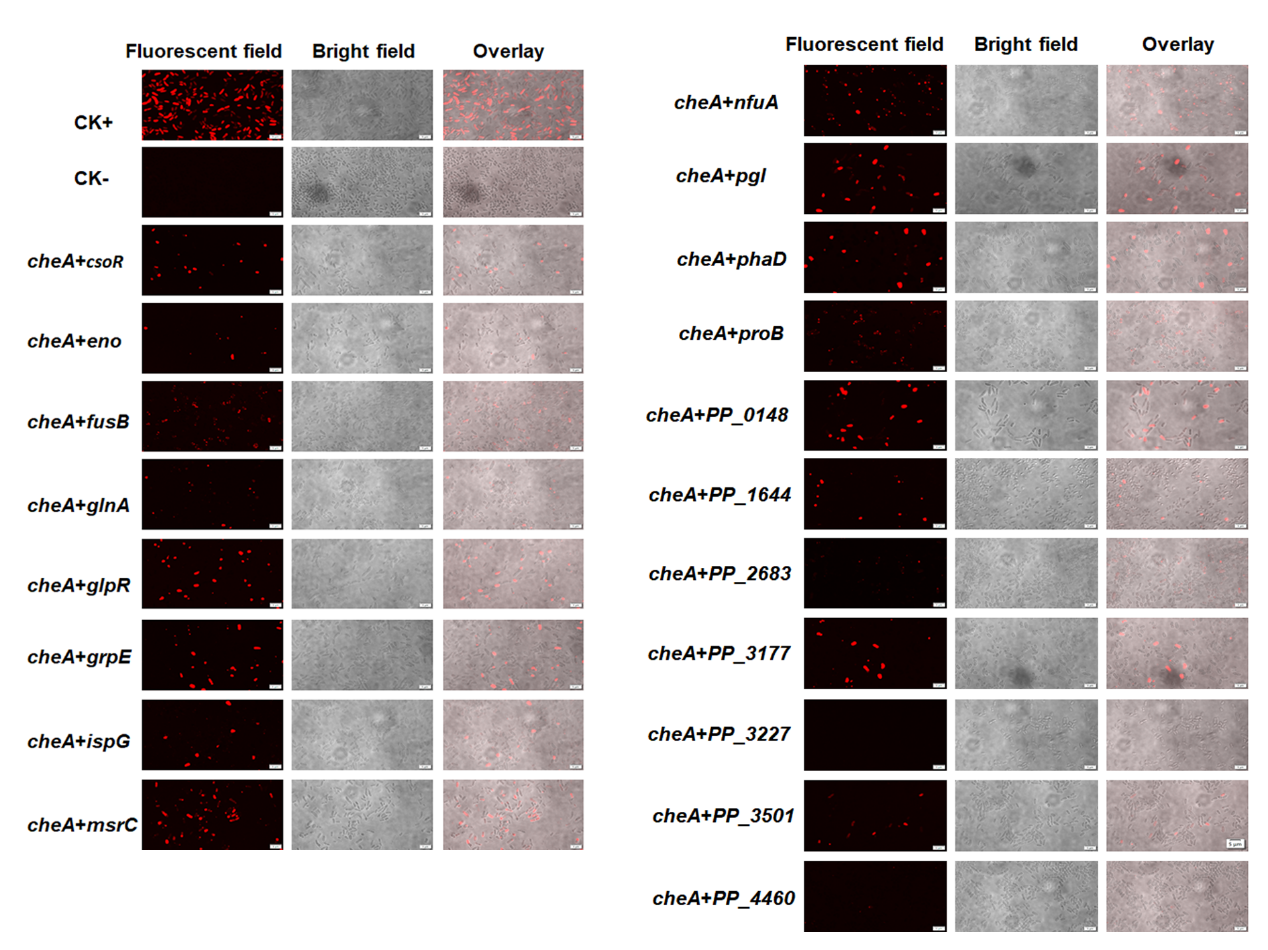

**Fig. S2** Detect the interaction between CheA and indicated proteins using BiFC. Representative images for each pair of proteins were shown. Jun-KN151+Fos-LC151 and KN151+LC151 were used as CK+ and CK-, respectively. RFP channel images (fluorescent field), bright field images, and overlay images of the same field are shown. Scale bar = 2 μm.

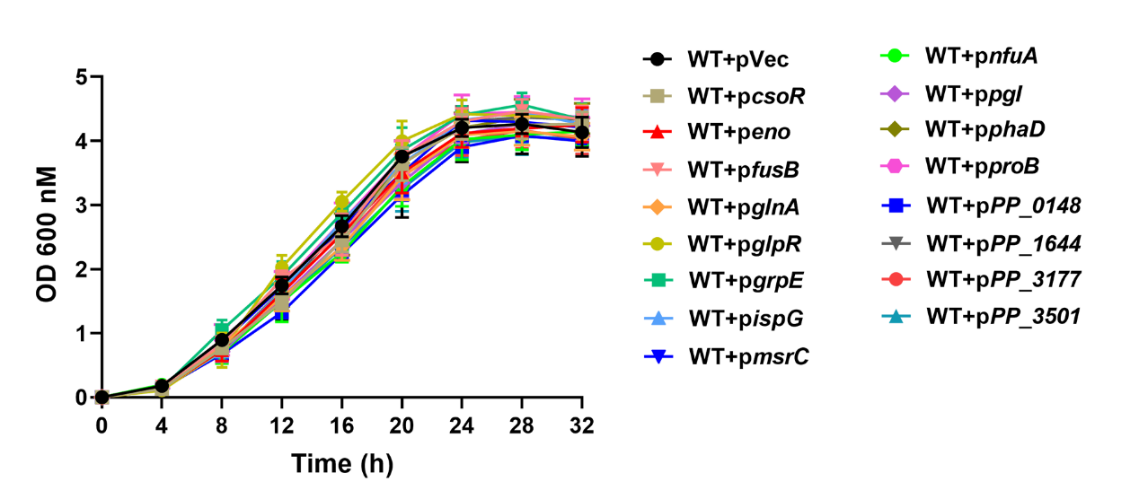

**Fig. S3** Growth curve of the 16 overexpression strains and wild-type strain in liquid LB broth (100 mL in a 250 mL triangular glass flask, at 28°C with 180 rpm shaking). The optical density at 600 nm (OD_600_) was used to characterize the growth of bacterial cells in the medium.

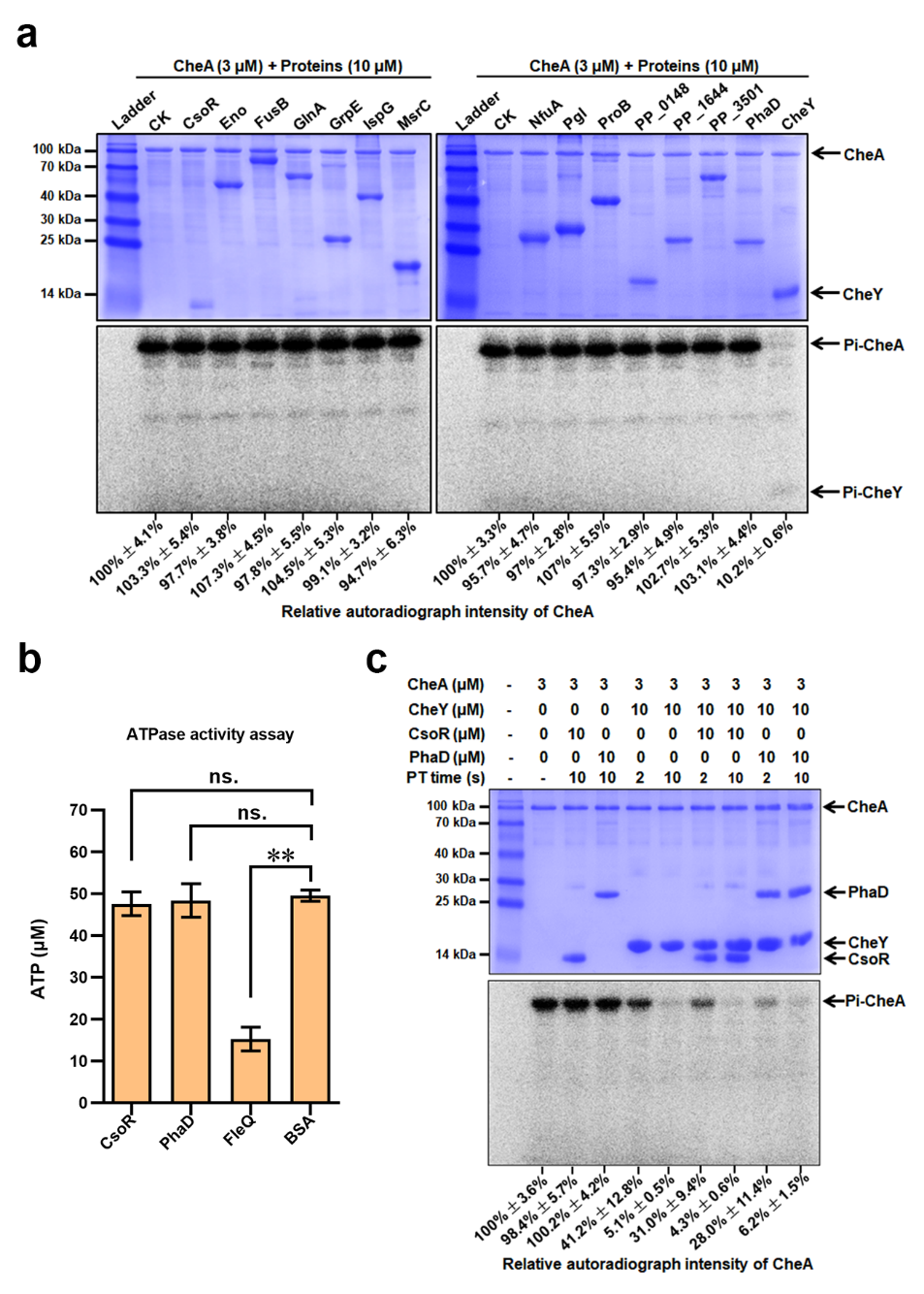

**Fig. S4 Role of target proteins in the CheA-mediated transphosphorylation.** (a) Transphosphorylation between CheA and the 14 proteins. Target proteins were added to the phosphorylated CheA and incubated for 10 s before adding termination buffer. CheY was added as a positive control. (b) ATPase activity of indicated proteins. The amount of remaining ATP after incubation with 10 μM indicated proteins for 30 min. BSA was used as a negative control. The asterisks represent statistically significant differences between FleQ and BSA (***P* < 0.01). “ns.” represents no statistically significant between the indicated protein and BSA. (c) Effect of CsoR and PhaD on the transphosphorylation between CheA and CheY. The PT time in seconds (2 s and 10 s) represents the time of transphosphorylation. The bands of CheA, CheY, CsoR, and PhaD on the gel were indicated with arrows. The relative autoradiograph intensity of the CheA band was calculated using Image J software, shown below each lane.

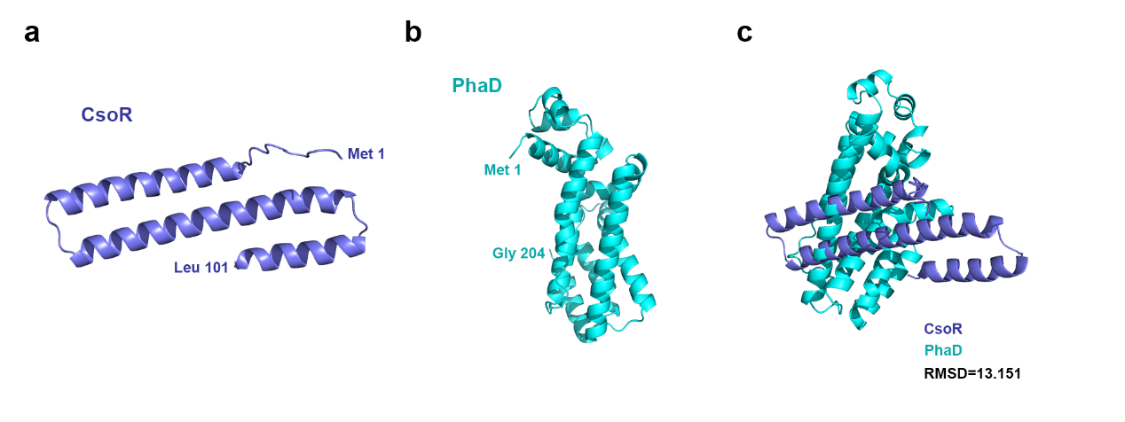

**Fig. S5** Predicted structure of the *P. putida* CsoR (a) and PhaD (b) obtained using an online AlphaFold server (https://www.alphafold.ebi.ac.uk). Position of the first and the last amino acid residue of CsoR and PhaD are indicated on the structure. (c) The structure alignment of CsoR and PhaD was performed with PyMOL software. The protein name and its structure were shown with the same color. The differences between protein structures were quantified using the Root Mean Square Deviation (RMSD) index. An RMSD value lower than 2 was considered a high similarity between two compared proteins.

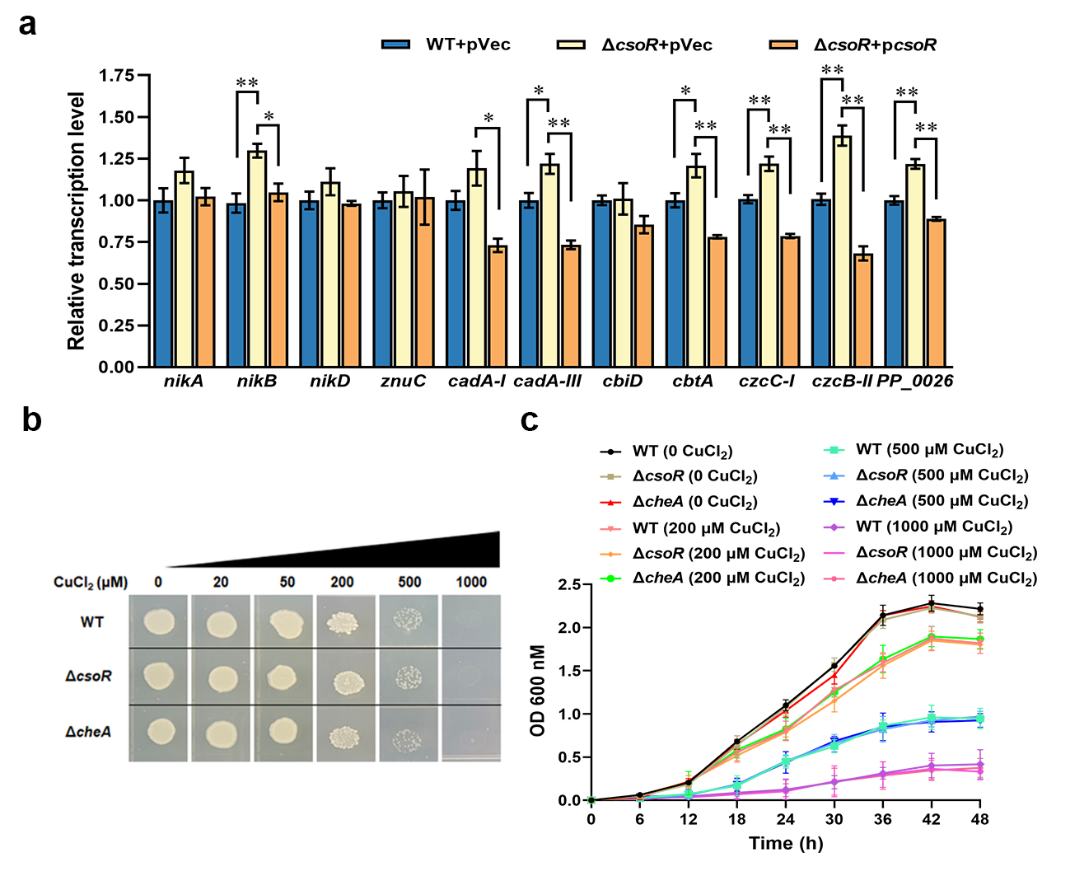

**Fig. S6** **Function of CsoR in the expression of metal resistant genes and the bacterial growth under copper stress.** (a) Analysis of relative transcription level of metal resistant genes in wild-type (WT+pVec), *csoR* mutant (Δ*csoR*+pVec), and complemented strain (Δ*csoR*+p*csoR*) by qRT-PCR. The results are the average of three independent assays. Error bars represent standard deviations. The asterisks represent statistically significant differences between the two compared strains (**P* < 0.05, ***P* < 0.01). (b and c) Effect of *csoR*/*cheA* deletion on bacterial growth under copper stress. Growth of WT, Δ*csoR*, and Δ*cheA* on M9 minimal medium agar plate (b) and in liquid 1/4 LB medium (c) containing different CuCl_2_ concentrations. A same amount of bacterial culture (OD600=0.5) was spotted onto the plate and incubated at 28°C for 24 h. The concentrations of CuCl_2_ used in the assay were indicated above.

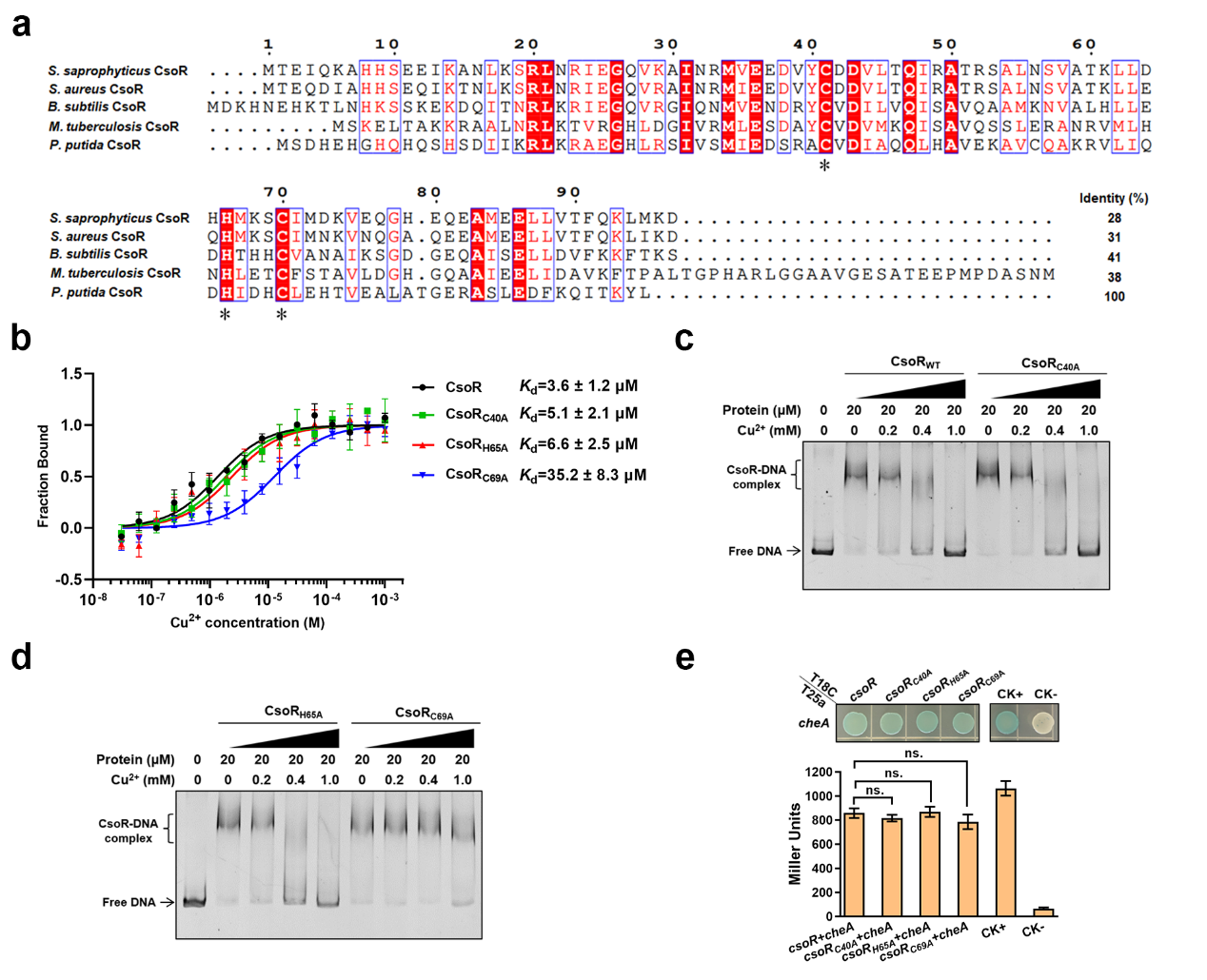

**Fig. S7 Role of the three conserved residues in the Cu^2+^-binding ability of CsoR.** (a) *P. putida* CsoR and CsoR homologs of indicated bacterial species are aligned using an online ClustalW server (https://www.genome.jp/tools-bin/clustalw). The number above the sequence represents the amino acid order of *S. saprophyticus* CsoR. The identical residues are shown as white on red letters. Similar residues are shown in red in blue boxes. Asterisks indicated the three conserved residues involved in Cu^2+^-binding. The sequence identity of CsoR homologs to the *P. putida* CsoR was calculated. (b) MST analysis of the interaction between wild-type and mutated CsoRs and Cu^2+^. Wild-type/mutated CsoR (250 nM) was incubated with increasing concentrations of Cu^2+^. (c and d) Effect of point mutation on the binding of CsoR to *copA-I* promoter in the presence of Cu^2+^ in the EMSA assay. Free DNA and CsoR-DNA complex are indicated. (e) Detect the interaction between CheA and point-mutated CsoR using BTH. The LacZ activities of colonies were shown below. Data are from three independent experiments, and error bars are the standard error of the means. “ns.” represents none statistically significant between indicated strain and CK- strain analyzed by Student’s t-test.

**Table S1** Target proteins identified in a pull-down assay. iBAQ_T (%) and iBAQ_CK (%) represent the percentage of certain proteins in experimental and control samples, respectively. The *P* values are for three technical replicates. Protein names and descriptions are based on the annotated genome as indicated at http://www.pseudomonas.com.

| Protein name/ID | iBAQ_T (%) | iBAQ_CK (%) | Log_2_ (iBAQ_T  /iBAQ _CK) | *P* value | Description |
| --- | --- | --- | --- | --- | --- |
| CheA | 14.0253 | 0.0582 | 7.9126 | 0.0014 | Chemotaxis histidine kinase |
| CheY | 1.3015 | 0.0133 | 6.6104 | 0.0009 | Chemotaxis response regulator |
| CheZ | 0.5521 | 0.0280 | 4.3030 | 0.0039 | Protein phosphatase |
| PP_2359 | 0.0128 | 0.0004 | 5.0712 | 0.0071 | Putative Type 1 pili subunit CsuA/B protein |
| PP_3104 | 0.0506 | 0.0018 | 4.7829 | 0.0046 | Hypothetical protein |
| SspB | 0.3662 | 0.0182 | 4.3305 | 0.0191 | ClpXP protease specificity-enhancing factor |
| PP_1958 | 0.0135 | 0.0007 | 4.2978 | 0.0040 | Hypothetical protein |
| PP_1644 | 0.0227 | 0.0014 | 4.0341 | 0.0173 | NAD(P)H dehydrogenase |
| Eno | 0.0121 | 0.0007 | 4.0245 | 0.0390 | Enolase |
| GrpE | 0.0237 | 0.0017 | 3.7815 | 0.0156 | Heat shock protein GrpE |
| PaxB | 0.0314 | 0.0023 | 3.7735 | 0.0464 | Toxin secretion ATP-binding protein |
| ProB | 0.0092 | 0.0007 | 3.7289 | 0.0249 | Glutamate 5-kinase |
| PP_2907 | 0.0157 | 0.0013 | 3.6138 | 0.0209 | DNA-binding response regulator |
| CsoR | 0.0144 | 0.0012 | 3.6097 | 0.0149 | metal-binding protein |
| PP_1274 | 0.0367 | 0.0034 | 3.4318 | 0.0010 | Oxidoreductase, short-chain dehydrogenase |
| PP_0148 | 0.0126 | 0.0012 | 3.3665 | 0.0241 | Hypothetical protein |
| SpuD | 0.0026 | 0.0003 | 3.3282 | 0.0186 | Spermidine/putrescine ABC transporter substrate-binding protein |
| IspG | 0.0112 | 0.0012 | 3.2524 | 0.0051 | 4-hydroxy-3-methylbut-2-en-1-yl diphosphate synthase |
| GlnA | 0.0511 | 0.0057 | 3.1544 | 0.0360 | Glutamine synthetase |
| PP_3227 | 0.0041 | 0.0005 | 3.1500 | 0.0256 | Transcriptional regulator, LysR family |
| PP_5006 | 0.0139 | 0.0016 | 3.1473 | 0.0435 | Transcriptional regulator, TetR family |
| PP_3177 | 0.0055 | 0.0006 | 3.1171 | 0.0086 | Hypothetical protein |
| PP_3501 | 0.0017 | 0.0002 | 2.9666 | 0.0070 | Transposase |
| HflD | 0.0218 | 0.0029 | 2.9092 | 0.0142 | High frequency lysogenization protein HflD homolog |
| Pgl | 0.0246 | 0.0034 | 2.8652 | 0.0086 | 6-phosphogluconolactonase |
| MucA | 0.0186 | 0.0027 | 2.7673 | 0.0373 | Sigma factor AlgU negative regulatory protein |
| PP_3637 | 0.0101 | 0.0015 | 2.7629 | 0.0399 | Putative Sulfonate ABC transporter, ATP-binding protein |
| QuiP | 0.0022 | 0.0003 | 2.7169 | 0.0462 | Acyl-homoserine lactone acylase |
| PP_0568 | 0.0133 | 0.0022 | 2.6265 | 0.0054 | Y1_Tnp domain-containing protein |
| PP_4468 | 0.0133 | 0.0022 | 2.6077 | 0.0372 | Transcriptional regulator, Cro/CI family |
| PP_4460 | 0.0126 | 0.0023 | 2.4519 | 0.0406 | Transcriptional regulator, LysR family |
| PP_3640 | 0.0190 | 0.0036 | 2.4184 | 0.0236 | Transcriptional regulator, AraC family |
| PP_0120 | 0.0200 | 0.0038 | 2.4081 | 0.0181 | Putative zinc ABC transporter |
| NfuA | 0.0102 | 0.0019 | 2.4026 | 0.0262 | Fe/S biogenesis protein |
| PP_1489 | 0.0735 | 0.0147 | 2.3171 | 0.0248 | CheW domain protein |
| GlpR | 0.0371 | 0.0075 | 2.3065 | 0.0257 | DNA-binding transcriptional repressor |
| DctA | 0.0146 | 0.0031 | 2.2384 | 0.0269 | C4-dicarboxylate transport protein |
| MsrC | 0.0123 | 0.0027 | 2.1800 | 0.0488 | Free methionine-(R)-sulfoxide reductase |
| TatA-II | 0.0127 | 0.0028 | 2.1656 | 0.0168 | Sec-independent protein translocase protein |
| Gpx | 0.0123 | 0.0030 | 2.0279 | 0.0025 | Glutathione peroxidase |
| PP_4932 | 0.0293 | 0.0067 | 2.1332 | 0.0414 | Putative D-arabinose 1-dehydrogenase |
| FusB | 0.0052 | 0.0011 | 2.2250 | 0.0322 | Elongation factor G 2 |
| PP_2683 | 0.0029 | 0.0006 | 2.2718 | 0.0178 | Putative sensory box histidine kinase/response regulator |

**Table S2** Strains and plasmids used in this work.

| Strain or plasmid | Relevant genotype and/or description | Source or reference |
| --- | --- | --- |
| *E. coli* strains |  |  |
| DH5a | λ-Φ80dlacZΔM15Δ(*lacZYA*-*argF*)*U196 recA1 endA1 hsdR17*(rK- mK -) *supE44 thi-1 gyrA relA1* | Invitrogen Corp |
| S17-1/λpir | *RP4-2*(Km::Tn7,Tc::Mu-1), *pro-82*, *LAMpir*, *recA1*, *endA1*, *thiE1*, *hsdR17*, *creC510*, host for *pir*-dependent plasmids | Invitrogen Corp |
| BL21(DE3) | F-, *ompT*, *hsdS*(*rBB*-*mB*－), *gal*, *dcm*(DE3) | Invitrogen Corp |
| BTH101 | F-, cya-*99*, *araD139*, *galE15*, *galK16*, *rpsL1*, *hsdR2*, *mcrA1*, *mcrB1*, host for bacterial two-hybrid assay (BTH) | Stratagene |
| *P. putida* strains |  |  |
| WT | Wild type KT2440 | Lab stock |
| WT+pVec | Wild type KT2440 harboring empty vector pBBR1MCS-5 | Lab stock |
| WT+p*cheA* | Wild type KT2440 harboring empty vector pBBR1MCS-5-*cheA* | Lab stock |
| WT+p*csoR* | Wild type KT2440 harboring empty vector pBBR1MCS-5-*csoR* | Lab stock |
| Δ*csoR* | Unmarked *csoR* (*PP_2969*) deletion mutant | This work |
| Δ*csoR*+pVec | Δ*csoR* strain harboring empty vector pBBR1MCS-5 | This work |
| Δ*csoR*+p*csoR* | Δ*csoR* strain complemented with pBBR1MCS-5-*csoR* | This work |
| Δ*cheA* | Unmarked *cheA* (*PP_4338*) deletion mutant | This work |
| Δ*cheA*+pVec | Δ*cheA* strain harboring empty vector pBBR1MCS-5 | This work |
| Δ*cheA*+p*cheA* | Δ*cheA* strain complemented with pBBR1MCS-5-*cheA* | This work |
| Δ*cheA*Δ*csoR* | Unmarked *cheA csoR* double deletion mutant | This work |
| Δ*cheA*Δ*csoR*+pVec | Δ*cheA*Δ*csoR* strain harboring empty vector pBBR1MCS-5 | This work |
| Δ*cheA*Δ*csoR*+p*cheA* | Δ*cheA*Δ*csoR* strain complemented with pBBR1MCS-5-*cheA* | This work |
| Δ*cheA*Δ*csoR*+p*csoR* | Δ*cheA*Δ*csoR* strain complemented with pBBR1MCS-5-*csoR* | This work |
| Δ*cheA*Δ*csoR*+p*cheA*-*csoR* | Δ*cheA*Δ*csoR* strain complemented with pBBR1MCS-5-*cheA-csoR* | This work |
| Plasmids |  |  |
| pBBR401 | Knockout vector, derived from pBBR1-MCS5, replication fragment replaced by ori R6K replication fragment | Lab stock |
| pBBR401-*cheA*UP-DW | pBBR401 containing up and down homologous region of *cheA* | This work |
| pBBR401-*csoR*UP-DW | pBBR401 containing up and down homologous region of *csoR* | This work |
| pBBR1MCS5 | Expression vector containing gene coding α-subunit of *β*-galactosidase, Gm^r^ | Lab stock |
| pBBR1MCS5-*cheA* | pBBR1MCS5 containing intact *cheA* gene | This work |
| pBBR1MCS5-*cheA*_ΔHPT_ | pBBR1MCS5 containing truncated *cheA* without the region encoding HPT domain | This work |
| pBBR1MCS5-*cheA*_ΔYB_ | pBBR1MCS5 containing truncated *cheA* without the region encoding CheY binding domain | This work |
| pBBR1MCS5-*cheA*_ΔDim_ | pBBR1MCS5 containing truncated *cheA* without the region encoding dimerization domain | This work |
| pBBR1MCS5-*cheA*_ΔHATPase_ | pBBR1MCS5 containing truncated *cheA* without the region encoding HATPase domain | This work |
| pBBR1MCS5-*cheA*_ΔWB_ | pBBR1MCS5 containing truncated *cheA* without the region encoding CheW binding domain | This work |
| pBBR1MCS5-*csoR* | pBBR1MCS5 containing intact *csoR* gene | This work |
| pBBR1MCS5-*PP_5006* | pBBR1MCS5 containing intact *PP_5006* gene | This work |
| pET-28a | T7/his-tag expression vector, Km^r^ | Addgene |
| pHS-Strep | Protein expression vector with Strep II-tag, Km^r^ | This work |
| pKT-25a | Plasmid for BTH, Km^r^ | Lab stock |
| pKT-25a-*zip* | pKT-25a harboring *zip* gene, positive control in BTH | Lab stock |
| pUT-18C | Plasmid for bacterial two-hybrid, Amp^r^ | Lab stock |
| pUT-18C-*zip* | pUT-18C harboring *zip* gene, positive control in BTH | Lab stock |
| pBBR403-KN151-LC151 | Plasmid for bimolecular fluorescence complementation (BiFC), Gm^r^ | This work |
| pBBR403-Jun-KN151-Fos-LC151 | pBBR403-KN151-LC151 harboring Jun and Fos genes, positive control in BiFC | This work |

**Table S3** Primers used in this work.

| Primers | Sequence^a^ |
| --- | --- |
| 28a-CheA s | TGGGTCGCGGATCCATGAGCTTCGGCGCCG |
| 28a-CheA a | TGGTGGTGCTCGAGCCACCACCAGGACCTTGA |
| T18/T25-*cheA* s | CTAGTGGTGAATTCATGAGCTTCGGCGCCG |
| T18/T25-*cheA* a | ATCTAGATCTCGAGCCACCACCAGGACCTTGA |
| T18/T25-*csoR* s | CTAGTGGTGAATTCATGAGCGATCACGAACACG |
| T18/T25-*csoR* a | ATCTAGATCTCGAGCAGAACAGCCTGCTTCACC |
| T18/T25-*PP_1188* s | CTAGTGGTGAATTCATGACGACACGTCAGCCG |
| T18/T25-*PP_1188* a | ATCTAGATCTCGAGGGTGAAGTTCTGGGATGTGC |
| T18/T25-*PP_1612* s | CTAGTGGTGAATTCATGGCAAAAATCGTCGAC |
| T18/T25-*PP_1612* a | ATCTAGATCTCGAGGCAAGAGCAGAACGAGGA |
| T18/T25-*PP_4111* s | CTAGTGGTGAATTCACAACGCCCATCGAGCTG |
| T18/T25-*PP_4111* a | ATCTAGATCTCGAGGCGGTGCAAGAAGCTGTTG |
| T18/T25-*PP_5046* s | CTAGTGGTGAATTCATGTCGAAGTCGGTTCAAC |
| T18/T25-*PP_5046* a | ATCTAGATCTCGAGATAGAGCATAGCGTGACAGG |
| T18/T25-*PP_1074* s | CTAGTGGTGAATTCGGACCGCCCATGAATCTG |
| T18/T25-*PP_1074* a | ATCTAGATCTCGAGGATTACGGCAAGGTCATAGCAG |
| T18/T25-*PP_1874* s | CTAGTGGTGAATTCATGGCTGCAACGATGCTG |
| T18/T25-*PP_1874* a | ATCTAGATCTCGAGCGACCTGGTGCTGATGGACT |
| T18/T25-*PP_4728* s | CTAGTGGTGAATTCGATGAGCAGCTGAACGAGA |
| T18/T25-*PP_4728* a | ATCTAGATCTCGAGCAGAATGGAGACGCACGA |
| T18/T25-*PP_4015* s | CTAGTGGTGAATTCCGCATGAGCAACCTGCAG |
| T18/T25-*PP_4015* a | ATCTAGATCTCGAGACCGCAGTGAGCGAAGAAA |
| T18/T25-*PP_0853* s | CTAGTGGTGAATTCATGCACGGCGAATCTCC |
| T18/T25-*PP_0853* a | ATCTAGATCTCGAGCGACTGTTCTGGCAGGATG |
| T18/T25-*PP_1877* s | CTAGTGGTGAATTCATGATCGACCTCAATGCCA |
| T18/T25-*PP_1877* a | ATCTAGATCTCGAGGCCGAAGGTGTATGAAGAGC |
| T18/T25-*PP_1428* s | CTAGTGGTGAATTCATGAGTCGTGAAGCTTTGCA |
| T18/T25-*PP_1428* a | ATCTAGATCTCGAGCTGCCGTTGCGTTCGTAG |
| T18/T25-*PP_2378* s | CTAGTGGTGAATTCATGAGCGCTATAACCATTACC |
| T18/T25-*PP_2378* a | ATCTAGATCTCGAGGAAACCATCCTTTACAGGGA |
| T18/T25-*PP_0167* s | CTAGTGGTGAATTCGTGGAATCCGAAGTCAGTC |
| T18/T25-*PP_0167* a | ATCTAGATCTCGAGCCCATACAATCAAGAACACG |
| T18/T25-*PP_1023* s | CTAGTGGTGAATTCATGGGAGGGCGTGGTATG |
| T18/T25-*PP_1023* a | ATCTAGATCTCGAGCCTTGAGGCCATGCTGTGA |
| T18/T25-*PP_0691* s | CTAGTGGTGAATTCATGCGAAGCAAGGTGACG |
| T18/T25-*PP_0691* a | ATCTAGATCTCGAGGAACTTGAACACGGTGGACA |
| T18/T25-*PP_1108* s | CTAGTGGTGAATTCTTTCCTCCCTTCCGCCT |
| T18/T25-*PP_1108* a | ATCTAGATCTCGAGTGGACGTTAGTTCGGATGG |
| T18/T25-*PP_5181* s | CTAGTGGTGAATTCATGAATAAAATGGGCAAGAC |
| T18/T25-*PP_5181* a | ATCTAGATCTCGAGTGCCGCAATCATCAATTA |
| T18/T25-*PP_1321* s | CTAGTGGTGAATTCAGCTTTAAGGAGCCGTTG |
| T18/T25-*PP_1321* a | ATCTAGATCTCGAGCGTCAAGAAGCCAACTCG |
| T18/T25-*PP_5016* s | CTAGTGGTGAATTCATGGGTATCTTTGACTGGAA |
| T18/T25-*PP_5016* a | ATCTAGATCTCGAGGCTTCCTCTTCCATCTGC |
| T18/T25-*PP_0120* s | CTAGTGGTGAATTCGTGTCCCGATTCCTGGCT |
| T18/T25-*PP_0120* a | ATCTAGATCTCGAGCGACTGGTGCCTGGATGA |
| T18/T25-*PP_0148* s | CTAGTGGTGAATTCGTGATGCAGGATGTAACCG |
| T18/T25-*PP_0148* a | ATCTAGATCTCGAGCCAACGTGAAGCAGATCG |
| T18/T25-*PP_0568* s | CTAGTGGTGAATTCATGCAACGTCCAGGCTCC |
| T18/T25-*PP_0568* a | ATCTAGATCTCGAGTGGCGTTTCTTGGGTTCAT |
| T18/T25-*PP_1274* s | CTAGTGGTGAATTCATGCCCACCGTCCTGATC |
| T18/T25-*PP_1274* a | ATCTAGATCTCGAGACAGCCTGTACTGGCCTCTT |
| T18/T25-*PP_1489* s | CTAGTGGTGAATTCGCGATGAACGACTTGCAA |
| T18/T25-*PP_1489* a | ATCTAGATCTCGAGGCAGCCAGTAATCGTCCAG |
| T18/T25-*PP_1644* s | CTAGTGGTGAATTCGGAGATCCTGCCGTGAGC |
| T18/T25-*PP_1644* a | ATCTAGATCTCGAGCAGGTTGTTCACCACCAGCA |
| T18/T25-*PP_1958* s | CTAGTGGTGAATTCCCAGCGAGGCAGAGGGT |
| T18/T25-*PP_1958* a | ATCTAGATCTCGAGCTTGCCTGCTCCTCCTGAT |
| T18/T25-*PP_2359* s | CTAGTGGTGAATTCATGCGAACGAACCTTTCA |
| T18/T25-*PP_2359* a | ATCTAGATCTCGAGAACACGCTGACGCCAGT |
| T18/T25-*PP_2683* s | CTAGTGGTGAATTCATGGCGAGGCCCTCTGA |
| T18/T25-*PP_2683* a | ATCTAGATCTCGAGTGGCGTTACTCCATTGTCAGA |
| T18/T25-*PP_2907* s | CTAGTGGTGAATTCATGCCTCGCGTACTGACC |
| T18/T25-*PP_2907* a | ATCTAGATCTCGAGTGGAGACCTCGAAGTACAGCAC |
| T18/T25-*PP_3104* s | CTAGTGGTGAATTCGGGCAAGACGGCCTTCTC |
| T18/T25-*PP_3104* a | ATCTAGATCTCGAGGCCAATATCACGACCAGCAG |
| T18/T25-*PP_3177* s | CTAGTGGTGAATTCCCTGTCTTTGTGCTTTTGC |
| T18/T25-*PP_3177* a | ATCTAGATCTCGAGGAGGCTGTCCTTGAACGAGT |
| T18/T25-*PP_3227* s | CTAGTGGTGAATTCGATGATATGGACGCCTTCG |
| T18/T25-*PP_3227* a | ATCTAGATCTCGAGGCGTTGCCTAAGACCGTATT |
| T18/T25-*PP_3501* s | CTAGTGGTGAATTCATGACTTTCTCGCCCAATC |
| T18/T25-*PP_3501* a | ATCTAGATCTCGAGCAGTCAGAAGCGAGTTATGG |
| T18/T25-*PP_3637* s | CTAGTGGTGAATTCGTGTTGCTGGAGGCCCG |
| T18/T25-*PP_3637* a | ATCTAGATCTCGAGGCCGTTGTTATGTGCCACC |
| T18/T25-*PP_3640* s | CTAGTGGTGAATTCATGATTTCGTCCAGCCCG |
| T18/T25-*PP_3640* a | ATCTAGATCTCGAGGTGGCTATTCGGCTTGCTG |
| T18/T25-*PP_4460* s | CTAGTGGTGAATTCATGGAGAACAGAATCACCCT |
| T18/T25-*PP_4460* a | ATCTAGATCTCGAGGCCTTCCACTCGTATCAGC |
| T18/T25-*PP_4468* s | CTAGTGGTGAATTCATGCAGCTGAGAATTGCC |
| T18/T25-*PP_4468* a | ATCTAGATCTCGAGGCCGAACTCGCTTACCA |
| T18/T25-*PP_4932* s | CTAGTGGTGAATTCATGAGCCTGCCAACCCTG |
| T18/T25-*PP_4932* a | ATCTAGATCTCGAGTTGCGGATCAGCGTTCG |
| T18/T25-*PP_5006* s | CTAGTGGTGAATTCGACTGGATGAAAACCCGC |
| T18/T25-*PP_5006* a | ATCTAGATCTCGAGGCTGCAACACTACATCTCCAG |
| BIFC CheA s | AGGAAACAGAATTCATGCGTTTGATGAGCTTCGGC |
| BIFC CheA a | CTCGAGCCGAGCTCGGCGTAACGCTTGAGCAT |
| BIFC CsoR s | AAACTAGTGGATCCATGAGCGATCACGAACAC |
| BIFC CsoR a | CCGATCGCTCTAGAGAGGTACTTAGTGATTTGCTTG |
| BIFC PP_1612 s | AAACTAGTGGATCCATGGCAAAAATCGTCGAC |
| BIFC PP_1612 a | CCGATCGCTCTAGAGCGAAACTCGGCACGACC |
| BIFC PP_4111 s | AAACTAGTGGATCCCGCACAACGCCCATCGAG |
| BIFC PP_4111 a | CCGATCGCTCTAGACCCGCGGCTCTTCTTGAC |
| BIFC PP_5046 s | AAACTAGTGGATCCATGTCGAAGTCGGTTCAA |
| BIFC PP_5046 a | CCGATCGCTCTAGAGCAGCTGTAGTACAGCTC |
| BIFC PP_1074 s | AAACTAGTGGATCCATGAATCTGCCCCCCCGC |
| BIFC PP_1074 a | CCGATCGCTCTAGAGACCACCTCAAGCCTGAT |
| BIFC PP_4728 s | AAACTAGTGGATCCATGGCTGATGAGCAGCTG |
| BIFC PP_4728 a | CCGATCGCTCTAGAAGCCTTTTCATTGATCGA |
| BIFC PP_0853 s | AAACTAGTGGATCCATGCACGGCGAATCTCCG |
| BIFC PP_0853 a | CCGATCGCTCTAGAGCCACGAGCGATCAACGC |
| BIFC PP_1877 s | AAACTAGTGGATCCATGATCGACCTCAATGCC |
| BIFC PP_1877 a | CCGATCGCTCTAGAGCAGTCAGTCAACGCCAG |
| BIFC PP_2378 s | AAACTAGTGGATCCATGAGCGCTATAACCATT |
| BIFC PP_2378 a | CCGATCGCTCTAGAGTAGGCGTTTTCTTTCTG |
| BIFC PP_1023 s | AAACTAGTGGATCCATGGGAGGGCGTGGTATG |
| BIFC PP_1023 a | CCGATCGCTCTAGATGGGCACCAGTAGATGTC |
| BIFC PP_0691 s | AAACTAGTGGATCCATGCGAAGCAAGGTGACG |
| BIFC PP_0691 a | CCGATCGCTCTAGATACCAGCACCAGGTTGTC |
| BIFC PP_0148 s | AAACTAGTGGATCCGTGATGCAGGATGTAACCG |
| BIFC PP_0148 a | CCGATCGCTCTAGACAGGCGCTCCTCATCTTC |
| BIFC PP_1644 s | AAACTAGTGGATCCGCGCCCTACATCCTGGTG |
| BIFC PP_1644 a | CCGATCGCTCTAGACGCCGCCTCCAGGGCCTT |
| BIFC PP_2683 s | AAACTAGTGGATCCATGGCGAGGCCCTCTGAC |
| BIFC PP_2683 a | CCGATCGCTCTAGAATGCACCAGGAGCAACTG |
| BIFC PP_3177 s | AAACTAGTGGATCCGTGCTTTTGCACAGCGCT |
| BIFC PP_3177 a | CCGATCGCTCTAGAAGTTGGCAACAGCTGCAA |
| BIFC PP_3227 s | AAACTAGTGGATCCGATGATATGGACGCCTTC |
| BIFC PP_3227 a | CCGATCGCTCTAGAACCCAACAACCGATGTCC |
| BIFC PP_3501 s | AAACTAGTGGATCCATGACTTTCTCGCCCAAT |
| BIFC PP_3501 a | CCGATCGCTCTAGAGGCAGGCACCCACTGATG |
| BIFC PP_4460 s | AAACTAGTGGATCCATGGAGAACAGAATCACC |
| BIFC PP_4460 a | CCGATCGCTCTAGAGCTTCGGATTAGCAGATG |
| BIFC PP_5006 s | AAACTAGTGGATCCGACTGGATGAAAACCCGC |
| BIFC PP_5006 a | CCGATCGCTCTAGACCCCTCCAGGTACTTCAC |
| 403-CsoR s | AGGAAACAGAATTCATGAGCGATCACGAACACG |
| 403-CsoR a | ACTCTAGAGGATCCCAGAACAGCCTGCTTCACC |
| 403-PP_1612 s | AGGAAACAGAATTCATGGCAAAAATCGTCGAC |
| 403-PP_1612 a | ACTCTAGAGGATCCGCAAGAGCAGAACGAGGA |
| 403-PP_4111 s | AGGAAACAGAATTCGTCCACCCACAGGTAGTCC |
| 403-PP_4111 a | ACTCTAGAGGATCCGGAGCGGTGCAAGAAGC |
| 403-PP_5046 s | AGGAAACAGAATTCATGTCGAAGTCGGTTCAAC |
| 403-PP_5046 a | ACTCTAGAGGATCCATAGAGCATAGCGTGACAGG |
| 403-PP_1074 s | AGGAAACAGAATTCGGACCGCCCATGAATCTG |
| 403-PP_1074 a | ACTCTAGAGGATCCGATTACGGCAAGGTCATAGCAG |
| 403-PP_4728 s | AGGAAACAGAATTCCTCGCAGTCCACAAAA |
| 403-PP_4728 a | ACTCTAGAGGATCCCCCGAAACTTGAATTTG |
| 403-PP_0853 s | AGGAAACAGAATTCATGCACGGCGAATCTCC |
| 403-PP_0853 a | ACTCTAGAGGATCCCGACTGTTCTGGCAGGATG |
| 403-PP_1877 s | AGGAAACAGAATTCATGATCGACCTCAATGCCA |
| 403-PP_1877 a | ACTCTAGAGGATCCGCCGAAGGTGTATGAAGAGC |
| 403-PP_2378 s | AGGAAACAGAATTCATGAGCGCTATAACCATTACC |
| 403-PP_2378 a | ACTCTAGAGGATCCGAAACCATCCTTTACAGGGA |
| 403-PP_1023 s | AGGAAACAGAATTCATGGGAGGGCGTGGTATG |
| 403-PP_1023 a | ACTCTAGAGGATCCCCTTGAGGCCATGCTGTGA |
| 403-PP_0691 s | AGGAAACAGAATTCATGCGAAGCAAGGTGACG |
| 403-PP_0691 a | ACTCTAGAGGATCCGAACTTGAACACGGTGGACA |
| 403-PP_0148 s | AGGAAACAGAATTCGTGATGCAGGATGTAACCG |
| 403-PP_0148 a | ACTCTAGAGGATCCCCAACGTGAAGCAGATCG |
| 403-PP_1644 s | AGGAAACAGAATTCCCGCCGGAAAAGGTGC |
| 403-PP_1644 a | ACTCTAGAGGATCCGCGATCAGGCCGAAGA |
| 403-PP_3177 s | AGGAAACAGAATTCCCTGTCTTTGTGCTTTTGC |
| 403-PP_3177 a | ACTCTAGAGGATCCGAGGCTGTCCTTGAACGAGT |
| 403-PP_3501 s | AGGAAACAGAATTCATGACTTTCTCGCCCAATC |
| 403-PP_3501 a | ACTCTAGAGGATCCCAGTCAGAAGCGAGTTATGG |
| 403-PP_5006 s | AGGAAACAGAATTCACGTGCTGACCCGATGA |
| 403-PP_5006 a | ACTCTAGAGGATCCCCTACAAGATGCGCCAAA |
| 28a-CsoR s | TGGGTCGCGGATCCATGAGCGATCACGAACACG |
| 28a-CsoR a | TGGTGGTGCTCGAGCAGAACAGCCTGCTTCACC |
| 28a-PP_1612 s | TGGGTCGCGGATCCATGGCAAAAATCGTCGAC |
| 28a-PP_1612 a | TGGTGGTGCTCGAGGCAAGAGCAGAACGAGGA |
| 28a-PP_4111 s | TGGGTCGCGGATCCACAACGCCCATCGAGCTG |
| 28a-PP_4111 a | TGGTGGTGCTCGAGGCGGTGCAAGAAGCTGTTG |
| 28a-PP_5046 s | TGGGTCGCGGATCCATGTCGAAGTCGGTTCAAC |
| 28a-PP_5046 a | TGGTGGTGCTCGAGATAGAGCATAGCGTGACAGG |
| 28a-PP_1074 s | TGGGTCGCGGATCCGGACCGCCCATGAATCTG |
| 28a-PP_1074 a | TGGTGGTGCTCGAGGATTACGGCAAGGTCATAGCAG |
| 28a-PP_4728 s | TGGGTCGCGGATCCGATGAGCAGCTGAACGAGA |
| 28a-PP_4728 a | TGGTGGTGCTCGAGCAGAATGGAGACGCACGA |
| 28a-PP_0853 s | TGGGTCGCGGATCCATGCACGGCGAATCTCC |
| 28a-PP_0853 a | TGGTGGTGCTCGAGCGACTGTTCTGGCAGGATG |
| 28a-PP_1877 s | TGGGTCGCGGATCCATGATCGACCTCAATGCCA |
| 28a-PP_1877 a | TGGTGGTGCTCGAGGCCGAAGGTGTATGAAGAGC |
| 28a-PP_2378 s | TGGGTCGCGGATCCATGAGCGCTATAACCATTACC |
| 28a-PP_2378 a | TGGTGGTGCTCGAGGAAACCATCCTTTACAGGGA |
| 28a-PP_1023 s | TGGGTCGCGGATCCATGGGAGGGCGTGGTATG |
| 28a-PP_1023 a | TGGTGGTGCTCGAGCCTTGAGGCCATGCTGTGA |
| 28a-PP_0691 s | TGGGTCGCGGATCCATGCGAAGCAAGGTGACG |
| 28a-PP_0691 a | TGGTGGTGCTCGAGGAACTTGAACACGGTGGACA |
| 28a-PP_0148 s | TGGGTCGCGGATCCGTGATGCAGGATGTAACCG |
| 28a-PP_0148 a | TGGTGGTGCTCGAGCCAACGTGAAGCAGATCG |
| 28a-PP_1644 s | TGGGTCGCGGATCCGATCCTGCCGTGAGCGCG |
| 28a-PP_1644 a | TGGTGGTGCTCGAGCAGGTTGTTCACCACCAGCA |
| 28a-PP_2683 s | TGGGTCGCGGATCCTTCATGGCGAGGCCCTCT |
| 28a-PP_2683 a | TGGTGGTGCTCGAGTGGCGTTACTCCATTGTCAGA |
| 28a-PP_3177 s | TGGGTCGCGGATCCCCTGTCTTTGTGCTTTTGC |
| 28a-PP_3177 a | TGGTGGTGCTCGAGGAGGCTGTCCTTGAACGAGT |
| 28a-PP_3227 s | TGGGTCGCGGATCCGATGATATGGACGCCTTCG |
| 28a-PP_3227 a | TGGTGGTGCTCGAGGCGTTGCCTAAGACCGTATT |
| 28a-PP_3501 s | TGGGTCGCGGATCCATGACTTTCTCGCCCAATC |
| 28a-PP_3501 a | TGGTGGTGCTCGAGCAGTCAGAAGCGAGTTATGG |
| 28a-PP_4460 s | TGGGTCGCGGATCCATGGAGAACAGAATCACCCT |
| 28a-PP_4460 a | TGGTGGTGCTCGAGGCCTTCCACTCGTATCAGC |
| 28a-PP_5006 s | TGGGTCGCGGATCCGACTGGATGAAAACCCGC |
| 28a-PP_5006 a | TGGTGGTGCTCGAGGCTGCAACACTACATCTCCAG |
| T18-*cheA*_△domain_ s | CTAGTGGTGAATTCATGAGCTTCGGCGCCG |
| T18-*cheA*_△domain_ a | ATCTAGATCTCGAGCCACCACCAGGACCTTGA |
| T18-HPT^CheA^ s | CTAGTGGTGAATTCCGTTTGATGAGCTTCGGC |
| T18-HPT^CheA^ a | ATCTAGATCTCGAGTGTCACTTCGGCACGCT |
| T18-CheY Binding^CheA^ s | CTAGTGGTGAATTCAACAGCATGTTCGGCCAG |
| T18-CheY Binding^CheA^ a | ATCTAGATCTCGAGAGCGGGCTTGGCCACC |
| T18-Dimer^CheA^ s | CTAGTGGTGAATTCGAGAAGCACGCCGCCA |
| T18-Dimer^CheA^ a | ATCTAGATCTCGAGGCGGCCGAAGACCTTCT |
| T18-HATPase_c^CheA^ s | CTAGTGGTGAATTCACCGACCTTGACAAGAACCT |
| T18-HATPase_c^CheA^ a | ATCTAGATCTCGAGGTTGCCCAGCATCACCAT |
| T18-CheW Binding^CheA^ s | CTAGTGGTGAATTCGTCATCAAGGTGCCGCTG |
| T18-CheW Binding^CheA^ a | ATCTAGATCTCGAGCCGAAATCAAATACGCCG |
| 401-*cheA*_UP_ s | GGCCCCCCCTCGAGGAACTGAACCAGACCCGC |
| 401-*cheA*_UP_ a | GGCTGCAGGAATTCCCGAAGCTCATCAAACGT |
| 401-*cheA*_DW_ s | agcttcggGAATTCCGGCGTATTTGATTTCGG |
| 401-*cheA*_DW_ a | GAACTAGTGGATCCTGTGCTGGATCAACACGA |
| 401-*csoR*_UP_ s | GGCCCCCCCTCGAGCATTTGGCGGTCGGATTC |
| 401-*csoR*_UP_ a | GGCTGCAGGAATTCCGTGATCGCTCATGCCTT |
| 401-*csoR*_DW_ s | CGATCACGGAATTCTCACTAAGTACCTCTAGGCCTT |
| 401-*csoR*_DW_ a | GAACTAGTGGATCCGGGTTCGGGAATCTCGTC |
| MCS5-*csoR* s | GGCCCCCCCTCGAGACATGGTCGGCGGTCAG |
| MCS5-*csoR* a | GGCTGCAGGAATTCCCCAGCAGGATGGCACT |
| 402-*cheA* s | AGGAAACAGAATTCATGAGCTTCGGCGCCG |
| 402-*cheA* a | ACTCTAGAGGATCCCCACCACCAGGACCTTGA |
| 402-*csoR* s | AGGAAACAGAATTCATGAGCGATCACGAACAC |
| 402-*csoR* a | ACTCTAGAGGATCCCCAGCAGGATGGCACT |
| 402-*cheA*-*csoR* s | TCGAAAAAACTAGTATGAGCGATCACGAACAC |
| 402-*cheA*-*csoR* a | CACAGCGCGGCCGCCCAGCAGGATGGCACT |
| 402-*PP_5006* s | AGGAAACAGAATTCATGAAAACCCGCGATCG |
| 402-*PP_5006* a | ACTCTAGAGGATCCCGTCGGCTGCAACACTAC |
| 28a-GFPCheA s1 | GCGGCAGCCATATGATGGTGAGCAAGGGCGAG |
| 28a-GFPCheA a1 | GGCGAATTCGGATCCCTTGTACAGCTCGTCCATGC |
| 28a-GFPCheA s2 | GGATCCGAATTCGCCATGAGCTTCGGCGCCG |
| 28a-GFPCheA a2 | TGGTGGTGCTCGAGCCACCACCAGGACCTTGA |
| 28a-GFPCsoR s1 | GCGGCAGCCATATGATGGTGAGCAAGGGCGAG |
| 28a-GFPCsoR a1 | GGCGAATTCGGATCCCTTGTACAGCTCGTCCATGC |
| 28a-GFPCsoR s2 | GGATCCGAATTCGCCATGAGCGATCACGAACACG |
| 28a-GFPCsoR a2 | TGGTGGTGCTCGAGCCCAGCAGGATGGCACT |
| EMSA *copA-I*_pro_ s | TGTCAAGCTACCTGCTGAGCAGGATCGCCACAAGCA |
| EMSA *copA-I*_pro_ a | GCCGTAGGCAAACGCGTAG |
| Qpcr *copA-I* s | GCGTACAACTTCCACAAG |
| Qpcr *copA-I* a | TTGAGCAGGTAGGTGTAG |
| Qpcr *copA-II* s | CCAGCCTCAGAGACTAAC |
| Qpcr *copA-II* a | CAGATCCGCATAGGTCAG |
| Qpcr *copB-II* s | GTTGCAGCTTCTGTATGG |
| Qpcr *copB-II* a | GCCATTCTCTCCGATGAA |
| Qpcr *rpoD* s | CCTGATCCAGGAAGGCAACAT |
| Qpcr *rpoD* a | CAGGTGGCATAGGTCGAGAACT |
| *copA-I*_pro_-lacZ s | CTGATGCCGGTACCGCAGGATCGCCACAAGCA |
| *copA-I*_pro_-lacZ a | TTAGTCATCTGCAGGCCGTAGGCAAACGCGTAG |
| *copA-II*_pro_-lacZ s | CTGATGCCGGTACCCGATCAAAGCACTCGTAATG |
| *copA-II*_pro_-lacZ a | TTAGTCATCTGCAGTCTCGTGGTTTTGCTTTGC |
| *PP_0588*_pro_-lacZ s | CTGATGCCGGTACCGGCGATGCGGTATTTGTTG |
| *PP_0588*_pro_-lacZ a | TTAGTCATCTGCAGCATGCCTTGTACATTGAACACT |
| *csoR*_pro_-lacZ s | CTGATGCCGGTACCATGTGCGCCATTGTTGC |
| *csoR*_pro_-lacZ a | TTAGTCATCTGCAGGTGTTCGTGATCGCTCAT |
| 28a-CsoR_C40A_ s1 | TGGGTCGCGGATCCATGAGCGATCACGAACAC |
| 28a-CsoR_C40A_ a1 | GACGGCGGCCCGGCTGTCTTCGATCATGCTGAC |
| 28a-CsoR_C40A_ s2 | AGCCGGGCCGCCGTCGACATCGCTCAGCAACTG |
| 28a-CsoR_C40A_ a2 | TGGTGGTGCTCGAGCAGAACAGCCTGCTTCAC |
| 28a-CsoR_H65A_ s1 | TGGGTCGCGGATCCATGAGCGATCACGAACAC |
| 28a-CsoR_H65A_ a1 | GTGATCGATGGCGTCTTGGATAAGCACACGCTT |
| 28a-CsoR_H65A_ s2 | GACGCCATCGATCACTGCCTGGAACACACCGTC |
| 28a-CsoR_H65A_ a2 | TGGTGGTGCTCGAGCAGAACAGCCTGCTTCAC |
| 28a-CsoR_C69A_ s1 | TGGGTCGCGGATCCATGAGCGATCACGAACAC |
| 28a-CsoR_C69A_ a1 | CAGGGCGTGATCGATATGGTCTTGGATAAGCAC |
| 28a-CsoR_C69A_ s2 | ATCGATCACGCCCTGGAACACACCGTCGAAGC |
| 28a-CsoR_C69A_ a2 | TGGTGGTGCTCGAGCAGAACAGCCTGCTTCAC |
| 28a-GFPCsoR_C40A_ s1 | GCGGCAGCCATATGATGGTGAGCAAGGGCGAG |
| 28a-GFPCsoR_C40A_ a1 | GTCGACGGCGGCCCGGCTGTCTTCGATCATGCT |
| 28a-GFPCsoR_C40A_ s2 | CGGGCCGCCGTCGACATCGCTCAGCAACTGCAT |
| 28a-GFPCsoR_C40A_ a2 | TGGTGGTGCTCGAGCCCAGCAGGATGGCACT |
| 28a-GFPCsoR_H65A_ s1 | GCGGCAGCCATATGATGGTGAGCAAGGGCGAG |
| 28a-GFPCsoR_H65A_ a1 | ATCGATGGCGTCTTGGATAAGCACACGCTTGGC |
| 28a-GFPCsoR_H65A_ s2 | CAAGACGCCATCGATCACTGCCTGGAACACACC |
| 28a-GFPCsoR_H65A_ a2 | TGGTGGTGCTCGAGCCCAGCAGGATGGCACT |
| 28a-GFPCsoR_C69A_ s1 | GCGGCAGCCATATGATGGTGAGCAAGGGCGAG |
| 28a-GFPCsoR_C69A_ a1 | CAGGGCGTGATCGATATGGTCTTGGATAAGCAC |
| 28a-GFPCsoR_C69A_ s2 | ATCGATCACGCCCTGGAACACACCGTCGAAGCA |
| 28a-GFPCsoR_C69A_ a2 | TGGTGGTGCTCGAGCCCAGCAGGATGGCACT |
| T18-*csoR*_C40A/H65A/C69A_ s | CTAGTGGTGAATTCATGAGCGATCACGAACAC |
| T18-*csoR*_C40A/H65A/C69A_ a | ATCTAGATCTCGAGCAGAACAGCCTGCTTCAC |
| PHS-CheA(strep-tag II) s1 | GGATCCGAATTCGAGCTC |
| PHS-CheA(strep-tag II) a1 | GCTGCTGCCCATGGTATA |
| PHS-CheA(strep-tag II) s2 | CCATGGGCAGCAGCAGCCATATGAGCTTCGGC |
| PHS-CheA(strep-tag II) a2 | TCGAATTCGGATCCGGCGTAACGCTTGAGCAT |
